# Specific connectivity with Operculum 3 (OP3) brain region in acoustic trauma tinnitus: a seed-based resting state fMRI study

**DOI:** 10.1101/676429

**Authors:** Agnès Job, Anne Kavounoudias, Chloé Jaroszynski, Assia Jaillard, Chantal Delon-Martin

## Abstract

Tinnitus mechanisms remain poorly understood. Our previous functional MRI (fMRI) studies demonstrated an abnormal hyperactivity in the right parietal operculum 3 (OP3) in acoustic trauma tinnitus and during provoked phantom sound perceptions without hearing loss, which lead us to propose a new model of tinnitus. This new model is not directly linked with hearing loss and primary auditory cortex abnormalities, but with a proprioceptive disturbance related to middle-ear muscles. In the present study, a seed-based resting-state functional MRI method was used to explore the potential abnormal connectivity of this opercular region between an acoustic trauma tinnitus group presenting slight to mild tinnitus and a control group. Primary auditory cortex seeds were also explored because they were thought to be directly involved in tinnitus in most current models. In such a model, hearing loss and tinnitus handicap were confounding factors and were therefore regressed in our analysis. Between-groups comparisons showed a significant specific connectivity between the right OP3 seeds and the potential human homologue of the premotor ear-eye field (H-PEEF) bilaterally and the inferior parietal lobule (IPL) in the tinnitus group. Our findings suggest the existence of a simultaneous premotor ear-eye disturbance in tinnitus that could lift the veil on unexplained subclinical abnormalities in oculomotor tests found in tinnitus patients with normal vestibular responses. The present work confirms the involvement of the OP3 subregion in acoustic trauma tinnitus and provides some new clues to explain its putative mechanisms.

## INTRODUCTION

Tinnitus (*i.e*, ringing or whistling in the ears) is a symptom that emerges with a variety of diseases (Bagueley et al., 2013), leading to the definition of several types of tinnitus. Among the different types of tinnitus, acoustic trauma tinnitus was thought to be the direct result of cochlear injury and deafferentations inducing maladaptive changes in the central nervous system. However, this generally accepted model did not fit various situations (e.g. some people have hearing loss without tinnitus and *vice versa*) and moreover treatments based on this model to abolish tinnitus in humans have proved unsuccessful (Lehner et al., 2016; Vanneste et al., 2013; Elgoyhen et al., 2015). Contrarily to the current acoustic trauma tinnitus model, we showed in our previous fMRI studies that tinnitus may not come from hearing loss itself, but from a somatosensory disturbance, most probably involving middle ear muscle proprioceptive disturbance (Job et al., 2011; Job et al. 2016), leading to erroneous integration of proprioceptive signals (i.e, illusory sound) in the brain. Indeed, our previous study aimed at stimulating the middle ear proprioceptors by applying specific click vibration rates to the ear (Job et al. 2016), which resulted in a tinnitus-like perception in healthy subjects at a particular frequency. The neural correlates of this stimulation were found in a small brain region in the right operculum 3 (OP3) (Job et al., 2016). This region was also hyperactivated in the fMRI study comparing sound detection in a tinnitus group and a control group (Job et al., 2012). In many tinnitus studies it may have been overlooked, because MRI spatial smoothing may mask it by merging it with the primary auditory cortex.

It has been shown that disturbed muscle proprioceptive inputs using mechanical vibration send erroneous messages to the brain as a form of illusory perception (Roll et al., 2004; Kavounoudias et al., 2008). Similarly, illusory auditory perceptions could occur in the auditory system as long as such proprioceptors are present in the middle-ear muscles, a result that was demonstrated by Kierner (Kierner et al., 1999). In acoustic trauma, the physical stress generated in the whole auditory system, including the middle ear, is extreme (Job et al, 2016; Wojtczak et al, 2017) and could overwhelm physiologic adaptations.

In order to further investigate the existence of a specific network related to tinnitus percept, we hereby explore the connectivity of this OP3 region in acoustic trauma tinnitus, using a seed-based connectivity approach with resting state functional magnetic resonance imaging (rs-fMRI). Seed-based rs-fMRI provides a close view of specific functional connectivity from brain regions of interest (ROIs). It also allows for adjustment of the data to confounding factors. Here, in this context, hearing loss may be a confounding physiological factor and psychological disturbance related to tinnitus may be a second confounding factor. If our hypothesis is valid, specific connectivity related to the percept of tinnitus *per se* should be observed despite these factors.

Possible differential connectivity between the tinnitus group and the control group was investigated from the right OP3 tinnitus-related seed (Job et al, 2016) and from its nearest regions, the anterior and posterior regions situated in the cytoarchitectonically defined right OP3 using same-shaped boxes. The left OP3 seed, mirror of the right OP3 seed, was also investigated for connectivity to check for laterality. Lastly, we explored connectivity from primary auditory cortex ROIs (Heschl’s gyri) since the current acoustic trauma tinnitus model involves these areas.

## MATERIAL AND METHODS

### Participants

Nineteen chronic tinnitus male participants (mean age 42. 5 ± 12 years old) with acoustic trauma sequelae ≥ 6 months (mean duration 11.9 ± 9.6 years, median = 12 years) and nineteen male controls matched in age without tinnitus (mean age 42. 5 ± 11. 9 years) were included in the study. All tinnitus participants were recruited through medical unit physicians in military regiments, through general announcements in the regiments and through the Ear, Nose and Throat (ENT) department of military hospitals open to civilians. The age-matched and gender-matched controls without tinnitus were recruited through announcement on a specialised website for call of volunteers for scientific protocols and by word-of-mouth. Control participants should not have reported frequent or permanent tinnitus, but could have experienced once or occasionally tinnitus in their ears.

The study was approved by the Local Ethics Committee CPP Sud-est V, Ref: 10-CRSS-05 MS 14-52, and conducted in accordance with the Declaration of Helsinki. Informed written consent was obtained from all participants.

### Hearing levels (HL), tinnitus description and Tinnitus Handicap Inventory (THI)

Both tinnitus participants and matched controls underwent audiograms. We assessed hearing performances by tonal audiometry at 0.25, 0.5, 1, 2, 4, 8 kHz (Audioscan fx by Essilor©) the day of the MRI acquisitions. Electroacoustic calibration of the audiometer equipped with Beyer dynamics DT48 earphones, was performed in accordance with AFNOR French standard S3007. We used the automated Hughson-Westlake procedure, which consisted in assessing hearing thresholds by modifying sound intensity using 5 dB up and 10 dB down steps. Sound intensity levels were modified automatically according to the participant’s responses. Threshold was defined as the lowest level at which at least two responses out of three presentations were obtained on an ascending run. Hearing losses were expressed in dB HL.

Two tinnitus questionnaires were completed by tinnitus participants. The first was an interview form collecting general information about the participant’s tinnitus. The second questionnaire was the tinnitus handicap inventory (THI) questionnaire, used internationally to robustly assess the intensity/intrusiveness of tinnitus and its associated handicap (Newman et al., 1996). THI scores are divided into severity grades (i.e., slight (THI 0-16), mild (THI 18-36), moderate (THI 38-56), severe (THI 58-76), catastrophic (78-100) (McCombe et al., 2001). According to this grading, the participants’ tinnitus characteristics were slight to moderate (Table 1).

**Table 1:**
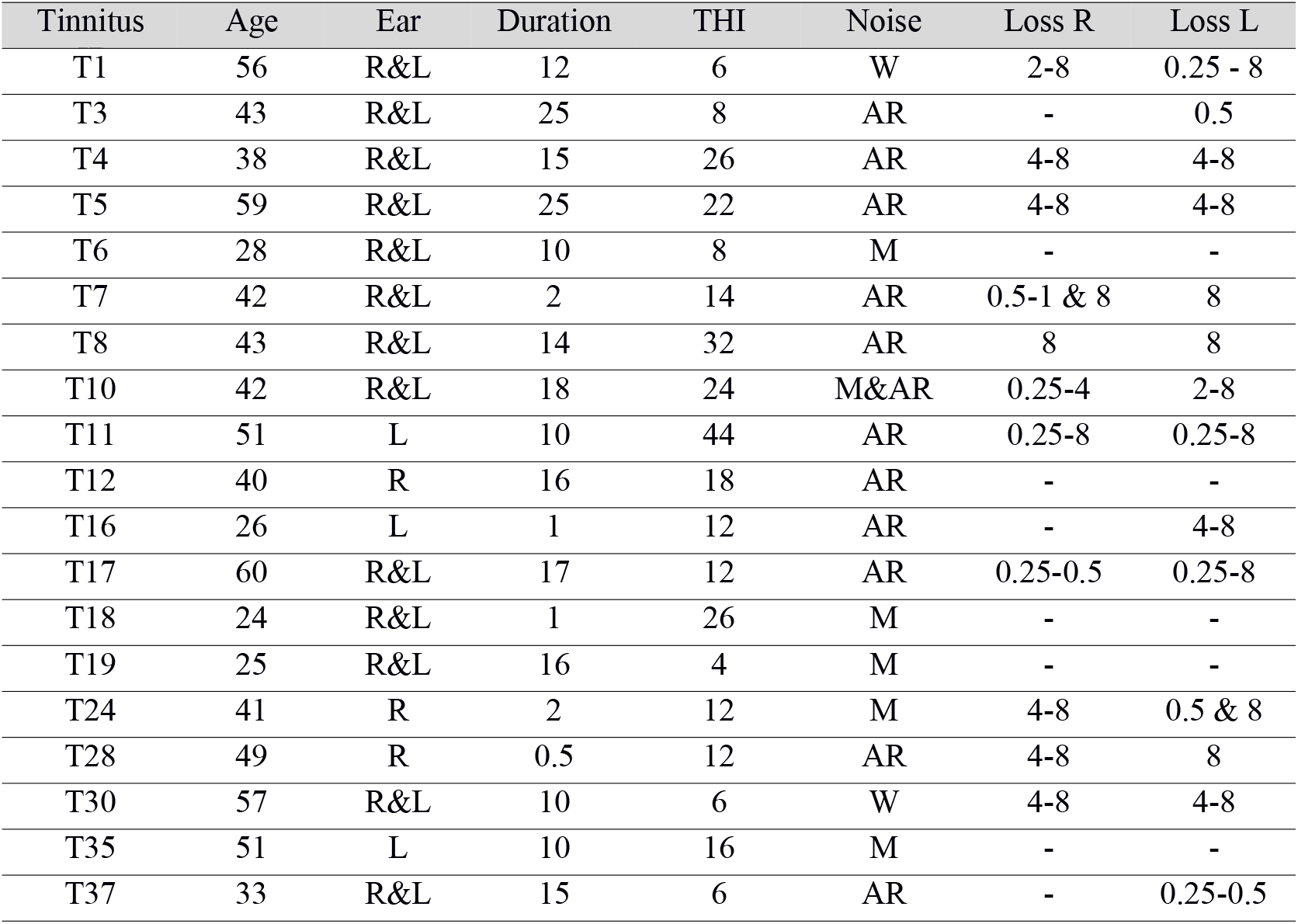
Tinnitus participant’s characteristics. Age (in years), Ear: Side of the tinnitus percept, Duration: tinnitus duration (in years), THI score: from Tinnitus Handicap Inventory questionnaire. Origin of the acoustic trauma: music (M), army riffles (AR), noisy workplace (W). Hearing Loss: frequency range (in kHz) where hearing loss > 25dB HL (hearing thresholds ≤ 25dB HL in frequencies tested are denoted (-)).

| Tinnitus | Age | Ear | Duration | THI | Noise | Loss R | Loss L |
| --- | --- | --- | --- | --- | --- | --- | --- |
| T1 | 56 | R&L | 12 | 6 | W | 2-8 | 0.25 - 8 |
| T3 | 43 | R&L | 25 | 8 | AR | - | 0.5 |
| T4 | 38 | R&L | 15 | 26 | AR | 4-8 | 4-8 |
| T5 | 59 | R&L | 25 | 22 | AR | 4-8 | 4-8 |
| T6 | 28 | R&L | 10 | 8 | M | - | - |
| T7 | 42 | R&L | 2 | 14 | AR | 0.5-1 & 8 | 8 |
| T8 | 43 | R&L | 14 | 32 | AR | 8 | 8 |
| T10 | 42 | R&L | 18 | 24 | M&AR | 0.25-4 | 2-8 |
| T11 | 51 | L | 10 | 44 | AR | 0.25-8 | 0.25-8 |
| T12 | 40 | R | 16 | 18 | AR | - | - |
| T16 | 26 | L | 1 | 12 | AR | - | 4-8 |
| T17 | 60 | R&L | 17 | 12 | AR | 0.25-0.5 | 0.25-8 |
| T18 | 24 | R&L | 1 | 26 | M | - | - |
| T19 | 25 | R&L | 16 | 4 | M | - | - |
| T24 | 41 | R | 2 | 12 | M | 4-8 | 0.5 & 8 |
| T28 | 49 | R | 0.5 | 12 | AR | 4-8 | 8 |
| T30 | 57 | R&L | 10 | 6 | W | 4-8 | 4-8 |
| T35 | 51 | L | 10 | 16 | M | - | - |
| T37 | 33 | R&L | 15 | 6 | AR | - | 0.25-0.5 |

### MRI data acquisition

A resting state-fMRI study was performed to search for a potential functional network specific to the tinnitus perception *per se*. For the rs-fMRI sequence, participants were instructed to lie quietly supine inside the scanner with their eyes open and to let their mind wanderi without focusing their thoughts on anything in particular. During acquisition, a grey background image with a small white cross in the centre was displayed. The MRI scanning sessions were conducted on a 3T Philips Achieva-TX scanner (Best, The Netherlands) at the Grenoble MRI facility-IRMaGe, equipped with a 32 channel-head coil. The examination protocol included a long rs-fMRI data sequence (longer than current sequence used in tinnitus research investigations) and a structural high resolution T1 weighted image. The parameters of the gradient echo EPI sequence were adjusted in order to lower the scanner noise to limit the acoustic dose received by participants. This was achieved by lowering the slope of the commutation gradients (i.e., SoftTone=yes). The frequency spectrum of the acquisition sequence presents a peak at a lower frequency (630 Hz) and with a 10 dB reduction as compared with the standard sequence. The high frequencies, above or equal to 1kHz, present an average reduction of 13 dB, when compared to the standard sequence used in fMRI on this MR scanner (Fig. 1). This sequence was reported as acoustically comfortable by participants. Other main parameters were: 32 slices, 3.5 mm-thick, acquired with a multiband factor of 2, in plane voxel size = 3×3 mm2, TR = 2000 ms, TE = 32 ms, flip angle = 75°, SENSE factor = 3, 5 dummy volumes and 400 volumes. This long acquisition duration (13’20”) was chosen since it is directly related to the reliability of rs-fMRI connectivity estimates (Birn et al., 2013; Termenon et al., 2016). This voxel size was chosen since it ensures a higher functional connectivity as compared to smaller size (Molloy et al, 2013). Choice of TR=2s avoids contamination of the useful part of the signal (below 0.1 Hz) by participant’s respiration. The examination ends with a T1-weighted structural image acquisition for each participant (3D MPRAGE, 0.9×0.9×1.2 mm^3^, TI = 800 ms, TR = 25 ms, TE = 3.9 ms, flip = 15°, SENSE factor = 2.2).

**Figure 1:**
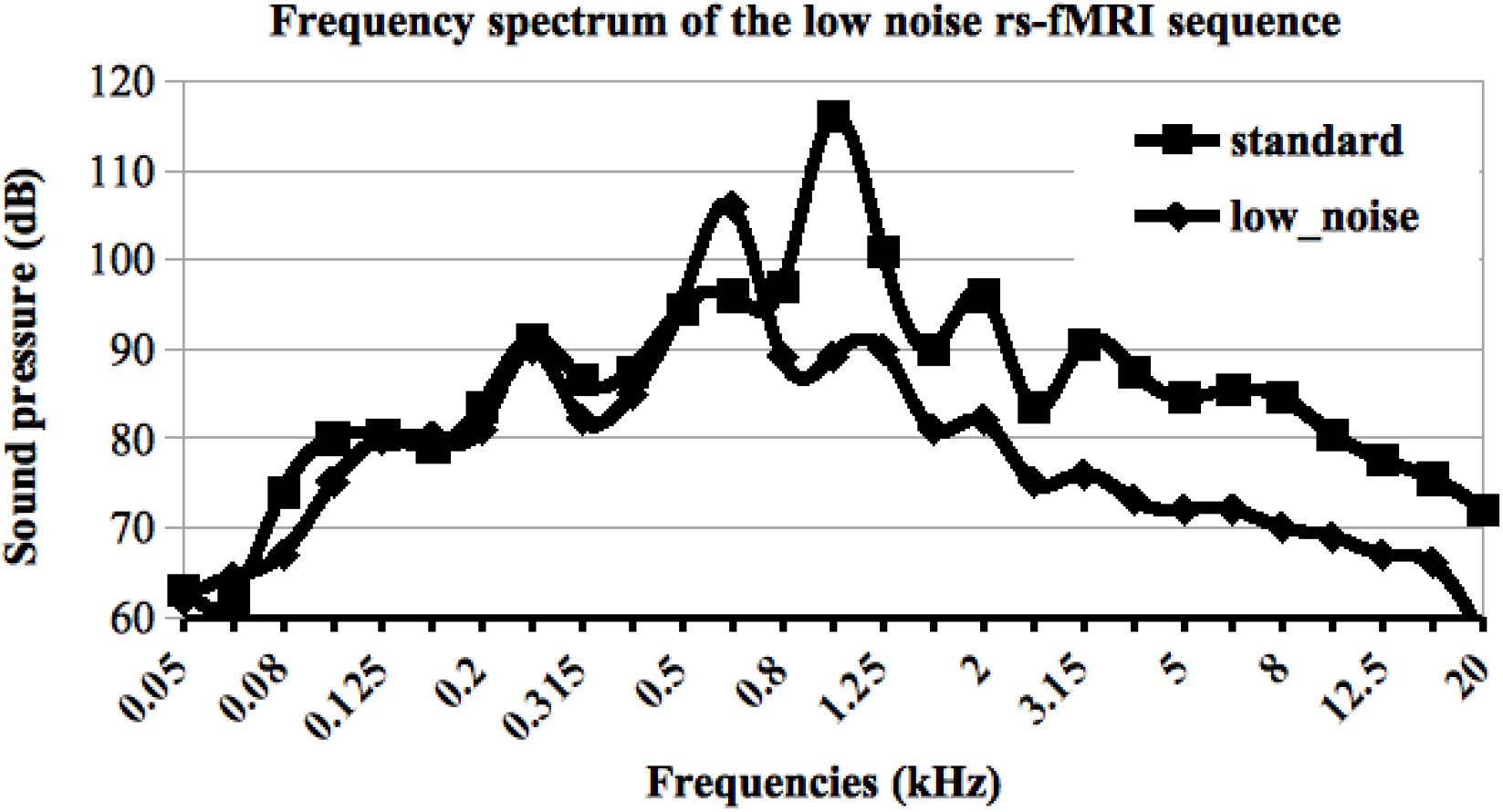
Frequency spectrum of the low noise resting state fMRI sequence used in this study (—) as compared to the frequency spectrum usually used with this scanner (---). The low noise sequence has been adjusted to limit high frequencies for all participants, with a mean attenuation of 13 dB above 1kHz.

### MRI data preprocessing

As the seed of interest in right OP3 is highly accurately located within the brain, we performed the analysis pipeline to preserve the spatial accuracy. We preprocessed the fMRI data using SPM12 software (www.fil.ion.ucl.ac.uk/spm/software/spm12/). Pre-processing of fMRI data included motion correction, slice-timing and co-registration of functional images on the individual structural image. No spatial smoothing was applied so as to avoid blurring of functional images. We further realigned all individual anatomical and functional images to a common referential using the DARTEL elastic registration algorithm (Ashburner, 2007) that allows accurate inter-individual brain realignment. To do this, we first performed segmentation of each structural image to extract the grey matter (GM), white matter (WM) and cerebrospinal fluid (CSF) images. We then calculated the deformation field that transforms the individual GM images to a high accurate GM template compatible with the standard MNI coordinates system. We chose the template provided with the Computational Anatomy Toolbox for SPM because it is spatially very accurate (http://dbm.neuro.uni-jena.de/cat). The deformation field image of each participant was further applied to the structural GM, WM, CSF images and to the functional individual images. All the individual images were then in the same template space with the same spatial resolution (isotropic 1.5 mm) for further accurate analysis. The mean structural image among participants was eventually computed for display purposes so that our results precisely matched the anatomy.

### Seed regions

We first included the tinnitus-related region of interest in right OP3, derived from our previous work (Job et al, 2016). This ROI and its spatial equivalent on the left hemisphere (as control) were boxes (9×4×4 mm^3^) built with Marsbar (http://marsbar.sourceforge.net), centered at MNI: ± 40 −13 17, corresponding to the location previously found. For an extensive exploration of right OP3, we created just anterior and posterior to the ROI, an anterior box centered at MNI: ± 40 −9 17 and a posterior box centered at MNI: ± 42 −17 17 (Fig. 2).

**Figure 2:**
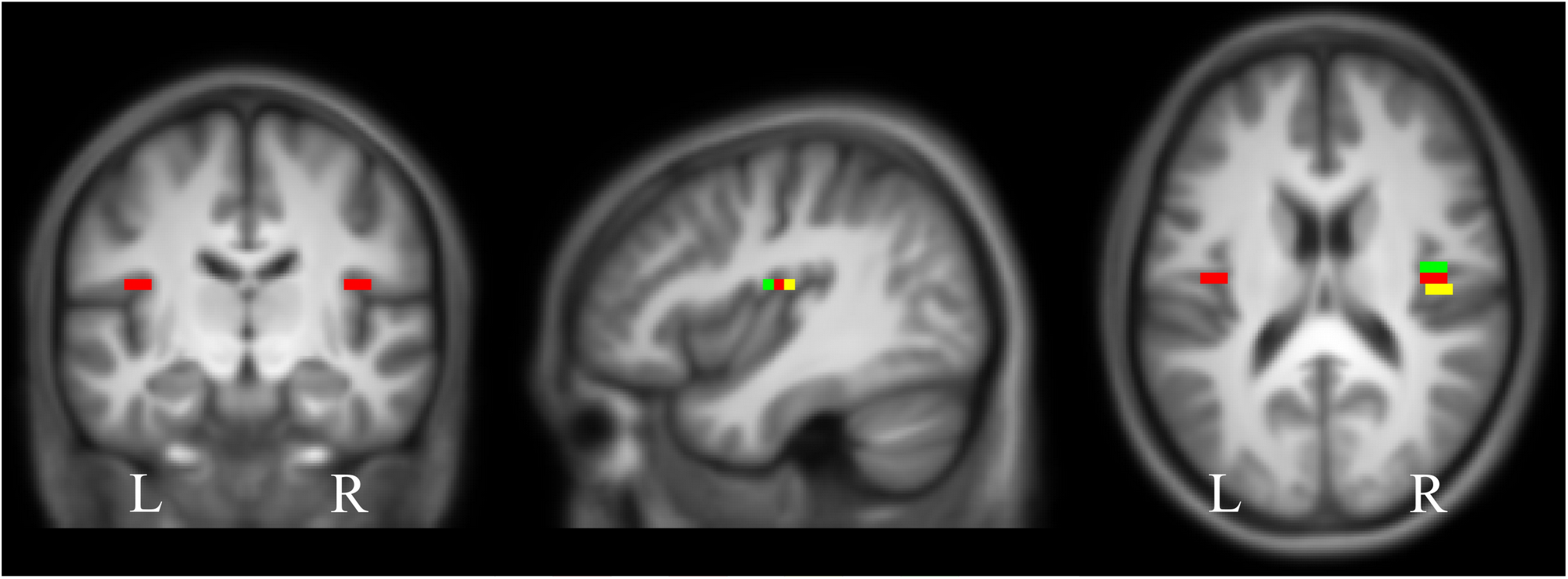
Seeds from OP3 region used in this connectivity study: the tinnitus-related seed (in red) (centered at MNI coordinates ±40, −13, 17) and the seeds from antero posterior gradient: the anterior seed (in green) and the posterior seed (in yellow). Localizations are superimposed on the mean anatomical images of our group of 38 subjects.

In order to further address the connectivity of the primary auditory areas that is said to be involved in tinnitus perception, two other seed regions were chosen in the right and left primary auditory cortices. These seeds were defined as the whole Heschl’s gyri by combining the Te1.0, Te1.1 and Te1.2 defined in the Anatomy toolbox (www.fz-juelich.de/inm/inm1/DE/Forschung/_docs/SPMAnatomyToolbox/SPMAnatomyToolbox_node.html).

### Functional seed-based connectivity analysis: individual level

We conducted seed-based connectivity analyses using the CONN toolbox (Whitfield-Gabrieli et al, 2012) (www.nitrc.org/projects/conn) version 18.a. For each individual, we first detected possible outliers in functional series derived from the art toolbox and then we removed physiological and other noise sources using the component-based noise correction method CompCor (Behzadi et al., 2007) that regresses signals from WM and CSF. The other components regressed are the six realignment parameters and their first-order temporal derivatives and outliers. No global signal regression is performed since its removal can affect correlation patterns and distort group differences (Saad et al., 2012). Prior to denoising, since the functional images were not spatially smoothed, the histogram was only weakly shifted towards the positive values. After this denoising step, the voxel-to-voxel correlation histogram was centred at about 0 with a standard deviation of about 0.12 for 107 degrees of freedom, which is a criterion of good data quality. This was visually checked for each participant’s data. Finally, bandpass filtering was applied within 0.008 to 0.08 Hz to all time-series, to select the data matching the resting-state frequency band. For each participant and each seed, Pearson’s correlation between time-series was computed voxel-wise, providing a correlation map that was eventually converted to z-map using Fisher’s r-to-z transform for further group analysis.

### Functional seed-based connectivity analysis: group level

Within- and between-group analyses from the control and the tinnitus groups were conducted for each seed separately. For the tinnitus group, the statistical tests were performed while regressing out potential confounds related to hearing loss and handicap related to tinnitus, by inserting the corresponding regressors of no interest (HL, THI). For within-group analysis, whole brain false discovery rate, FDR-corrected threshold of p<0.05 was retained for statistical significance. For between-group comparisons, to assess specific connectivity related to the tinnitus percept, a two sample t-test was performed. We analysed differential connectivity between the groups in the contrasts, tinnitus > controls and tinnitus < controls, from the different seeds to the entire brain. We chose a robust statistical threshold (p<0.0001, cluster-size p-FDR corrected < 0.05) to elicit significant differential connectivity.

## RESULTS

### Participants’ characteristics

In tinnitus participants, the mean hearing threshold was significantly different from that of the control group, for frequencies ranging from 1 kHz to 8 kHz (Mann-Whitney test, p<0.05), with marked hearing loss at frequencies superior to 4 kHz (Mann-Whitney test, p<0.001), as usually observed with acoustic traumas (Fig. 3). At the frequency of the peak of noise in the acquisition sequence, no differences in hearing were found between the two groups, thus ruling out the possibility that differential connectivity could arise from differences in acoustic perception related to scanner noise. All tinnitus participants described their tinnitus as high-pitched whistling except for one who described a medium high-pitched sizzling. The loudness was scored on an analogic visual scale (mean 4.5 ± sd 1.6 ranging from 2 to 8, from a total possible range 0-10). Bilateral tinnitus represented 68% of our population. There was a same amount of right and left tinnitus. Tinnitus duration ranged from 6 months to 25 years (mean 12.2 ± 7.3 years).

**Figure 3:**
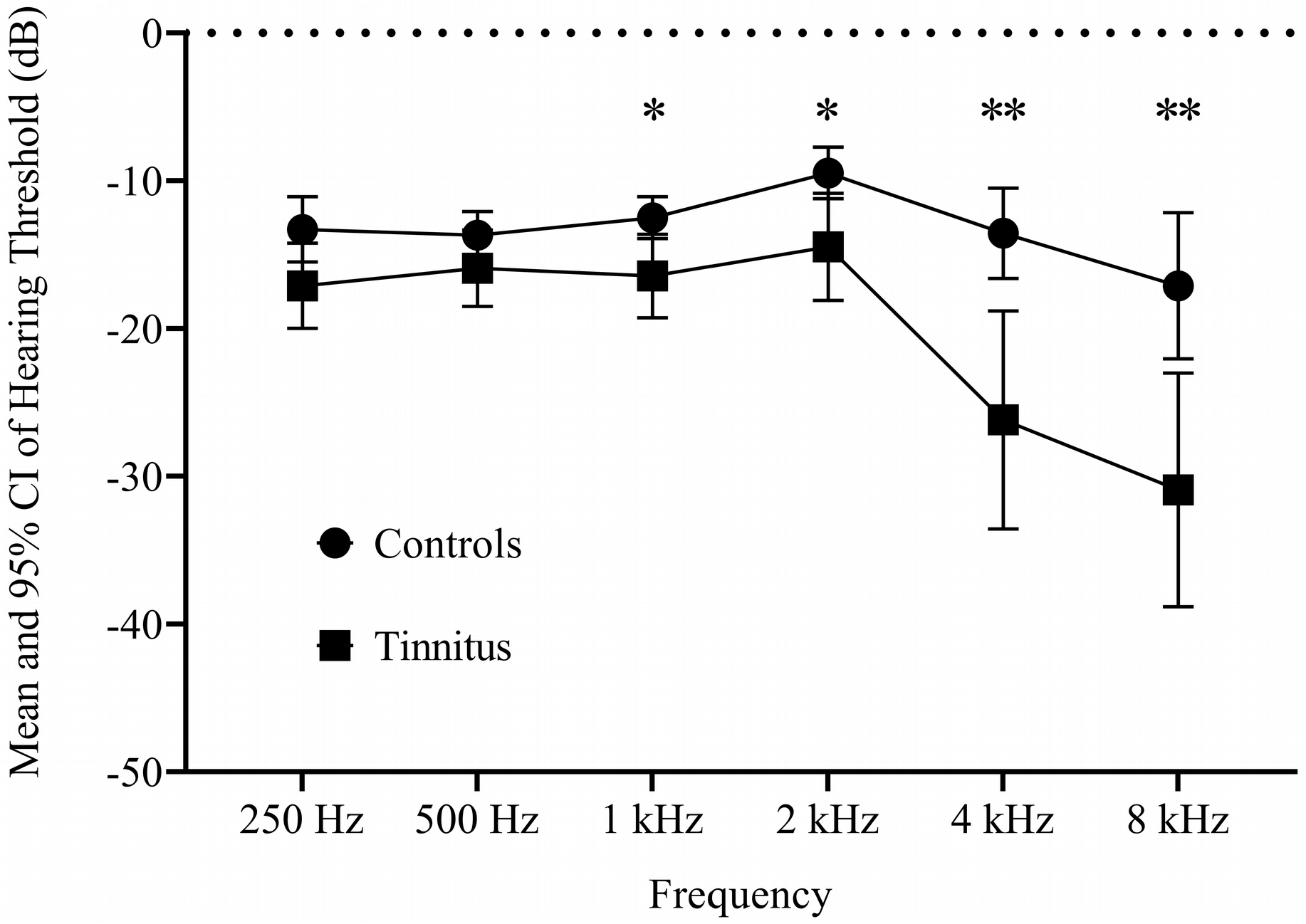
Audiograms of acoustic trauma tinnitus and control groups (Mean ± 95% CI) for both ears showing the hearing loss in high frequencies for the tinnitus group. Mann-Whitney tests results in the comparison of hearing thresholds (* p < 0.05, **p < 0.001).

Tinnitus Handicap Inventory scores described a slight to mild tinnitus in all participants (mean 16.2 ± 10.5) except for one with moderate tinnitus (THI=44), reflecting a tinnitus population less impacted than in the current literature in the field.

Detailed results from audiometry and neuropsychological testing are presented in Table 1 for tinnitus participants.

### Functional connectivity with OP3 seeds

Within-group connectivity of the right OP3 box in controls and tinnitus (Fig. 4A) revealed a main connectivity with the sensorimotor network (bilateral precentral gyri, bilateral postcentral gyri, bilateral central and parietal opercular cortices, supplementary motor area), but also with the salience network (bilateral insular cortex, anterior cingulate gyrus) and the auditory network (bilateral Heschl’s gyri, bilateral planum temporale, bilateral planum polare). Within-group connectivity of the contralateral left OP3 box in controls and tinnitus (Fig. 4B) provided a similar pattern as that of the right OP3 box, reflecting a symmetric pattern of connectivity. Note that for sake of visibility, the display threshold was lowered in the tinnitus group to make them visually comparable.

**Figure 4:**
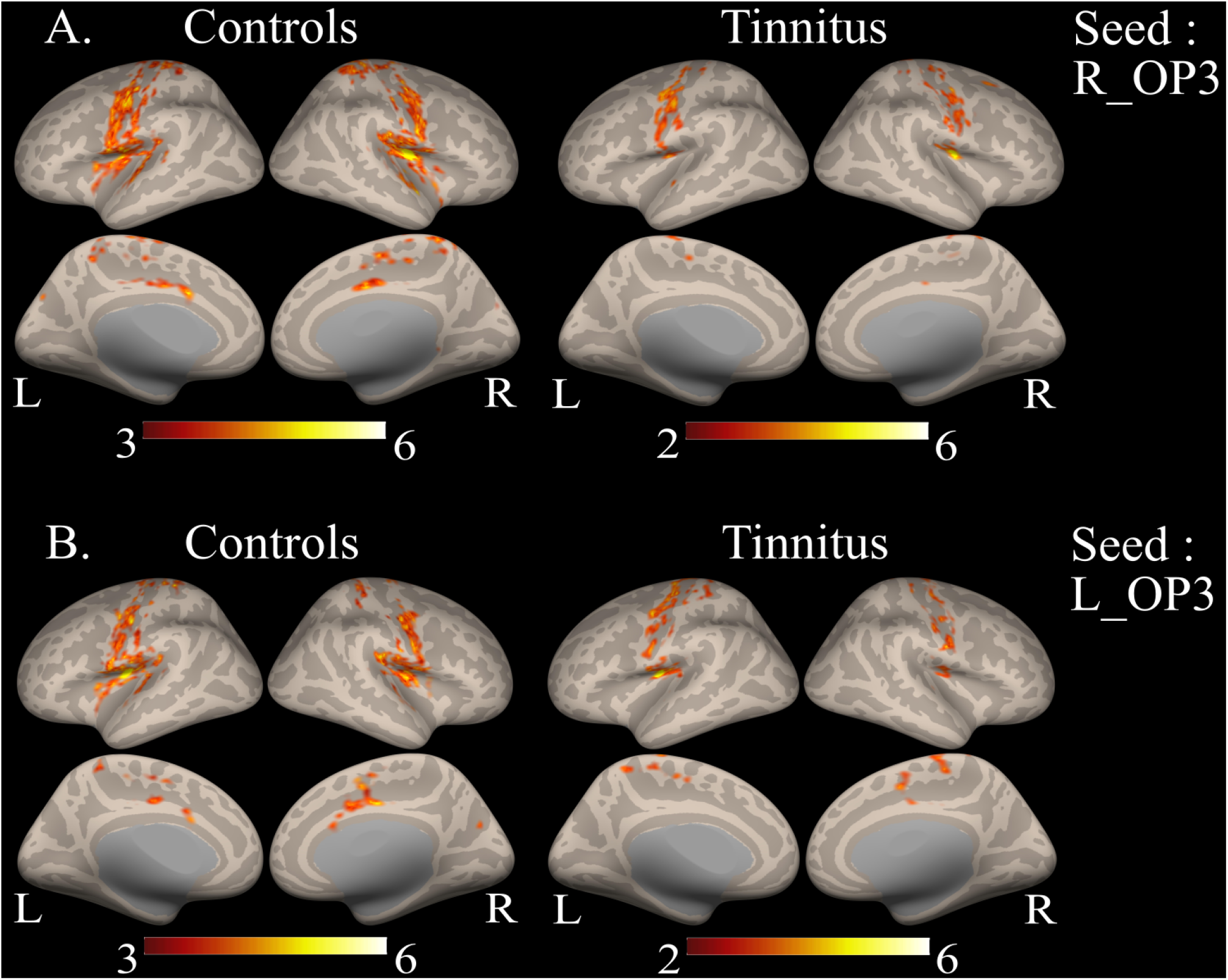
Within-group connectivity with the right (A) and left (B) OP3 seeds for controls and tinnitus groups showing a common pattern between groups and between left and right seeds. Results are superimposed on the MNI template provided with FreeSurfer.

Between-group comparison revealed a specific connectivity in the tinnitus group as compared to controls with the right OP3 seed. This specific connectivity was found with a region in the right superior frontal gyrus at the border of BA6 and 8 (Fig. 5A), with a maximum at MNI coordinates (20, 11, 47) (Table 2). We could not elicit any differential connectivity from the homologous left hemisphere OP3 box, even at a lower threshold. No significant reduced connectivity was found in tinnitus group as compared to the control group.

**Figure 5:**
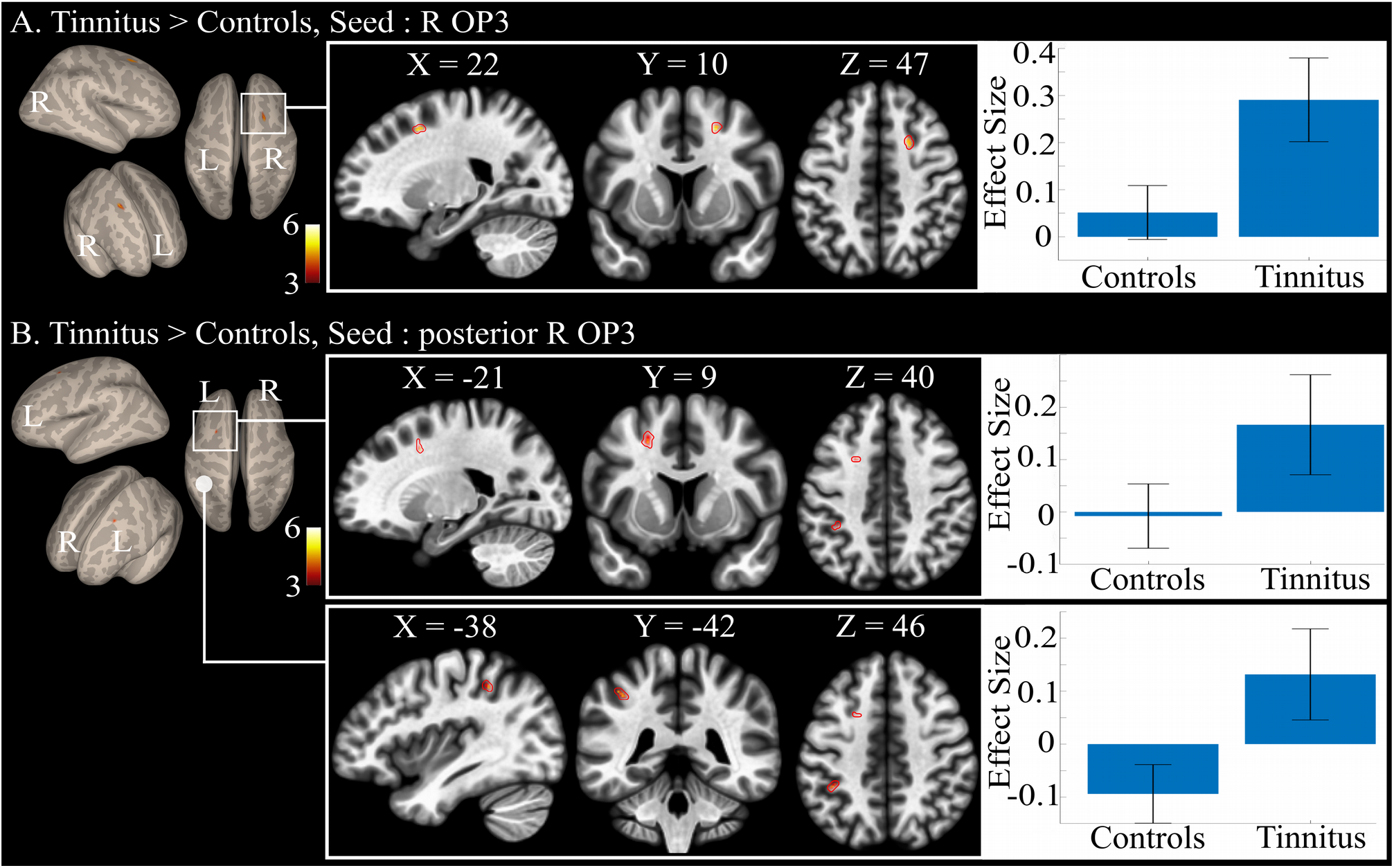
Between-group connectivity: (A) specific differential connectivity for tinnitus group as compared with the control group showing the increased connectivity of the right OP3 seed with the right superior frontal gyrus (R H-PEEF); (B) specific differential connectivity for tinnitus group as compared with the control group showing the increased connectivity of the right posterior OP3 seed with the left superior frontal gyrus (L H-PEEF) and left inferior parietal lobule. Results are superimposed on the MNI template provided with FreeSurfer.

**Table 2:** Significant differential functional connectivity results for Tinnitus>Controls groups for the right OP3 seed (in red, Fig. 2) and for the right OP3 posterior seed (in yellow, Fig. 2). No other significant differential connectivity was found between both groups. R: right, L: left. Results were all obtained at height threshold p< 0.0001, extent threshold pFDR corr<0.05. *: region achieving above height threshold but only a tendency for statistical extent threshold.

| TINNITUS > CONTROL |  |  | MNI coordinates | Statistical results |  |
| --- | --- | --- | --- | --- | --- |
| Seed | Brain region | BA | x, y, z | T-value | extent |
| R. OP3 box<br>(40, -13, 17) | R. superior frontal gyrus | 6/8 | 20, 11, 47 | 6.10 | 22 |
| R. OP3 post box<br>(42, -17, 17) | L. superior frontal gyrus | 6/8 | -21, 8, 44 | 5.17 | 9* |
|  | L. inferior parietal lobule | 40 | -42, -42, 48 | 5.12 | 18 |
| L. Heschl gyrus | Posterior cingulate cortex | 23 | 0, -47, 32 | 5.11 | 18 |

The anterior and posterior seeds in the right OP3 gave different between-group differential connectivity results. The anterior seed did not show any significant differential connectivity. In contrast, with the posterior right OP3 seed, we found a specific connectivity with the contralateral left H-PEEF and with the left inferior parietal lobule (IPL) in a part that could correspond to hIP2 or to PFt (Caspers et al, 2008), according to the Anatomy toolbox (Fig. 5B, Table 2).

### Functional connectivity with auditory seeds

Within-group connectivity with Heschl’s gyri showed bilateral connectivity with auditory regions, operculum, insula, dorsal anterior cingulate cortex and the face representation of the sensorimotor cortex (Fig. 6).

**Figure 6:**
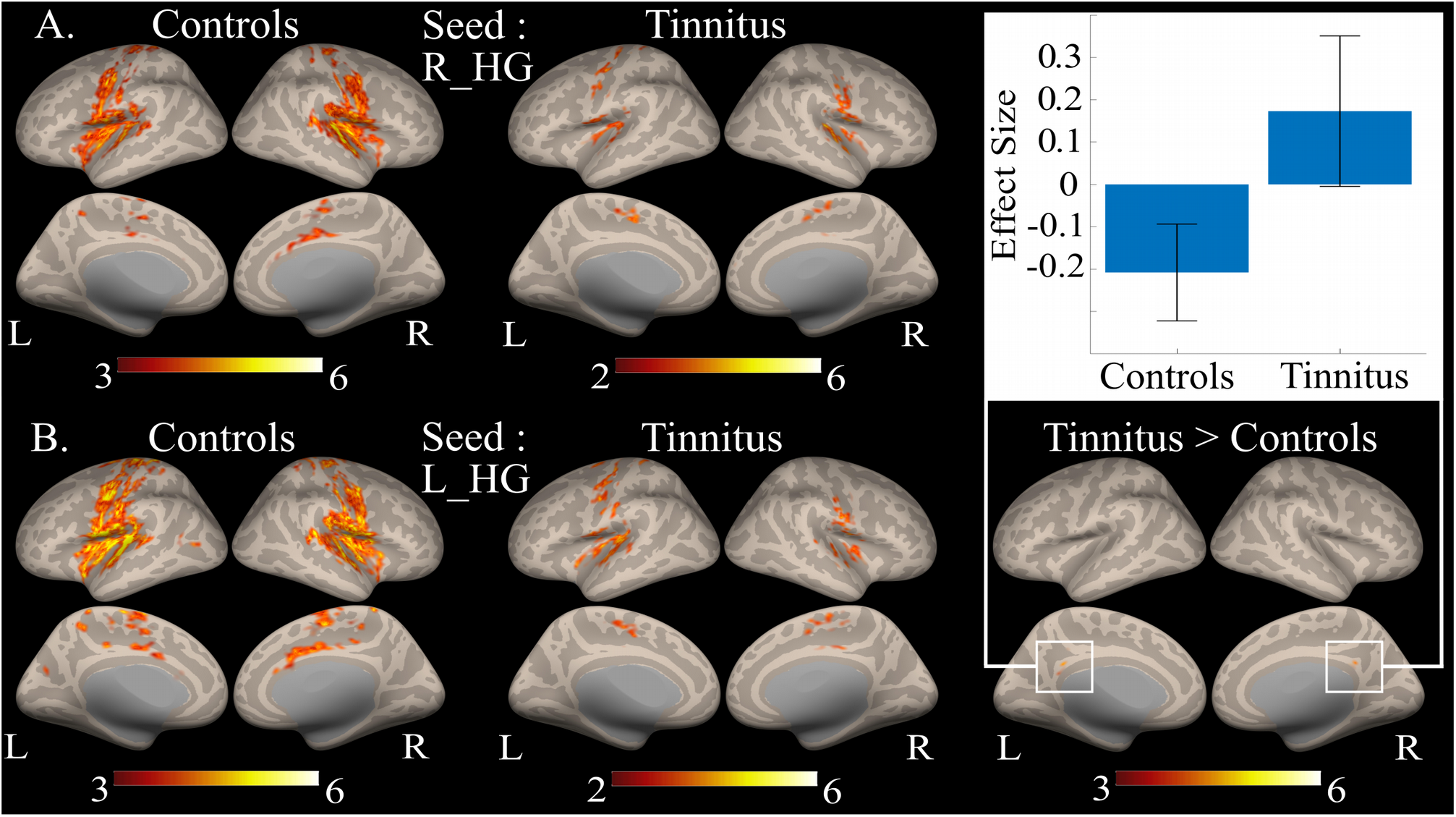
Within-group connectivity with the right (A) and left (B) Heschl’gyrus seeds for controls and tinnitus groups showing a common pattern between groups. Between-group differences reached significance for the single connectivity between left HG and the posterior cingulate cortex (right part of the figure). Results are superimposed on the MNI template provided with FreeSurfer.

Between-group connectivity with Heschl’s gyri seeds elicited a single significant differential connectivity between left Heschl’s gyrus and the bilateral posterior cingulate cortex. No other differential connectivity was found at the same statistical threshold than for the right OP3 seed, when adjusting for the hearing loss and for the handicap related to tinnitus.

## DISCUSSION

The tinnitus participants that were recruited in this study were different from those usually recruited. Indeed, in most studies, tinnitus subjects are recruited from ENT services and are strongly impacted psychologically by their tinnitus. The results found in the literature are thus related not only to tinnitus percept but also to psychological and quality of life factors and it is therefore difficult to disentangle these factors. On the contrary, in the present study, the tinnitus participants were not strongly affected by their tinnitus: it did not impact their quality of life even if they presented hearing loss at high frequencies and a slight handicap, as assessed by the THI score. In addition, the analyses that were performed here took these factors into account to regress their impact. Altogether, we thus believe that the results presented here mainly related to the tinnitus percept *per se*.

When considering the seed of interest, the right OP3 box derived from previous studies (Job et al. 2012, Job et al. 2016a), within-group connectivity was found to involve the somatosensory and the saliency as well as the auditory networks. This suggests that our seed may play a pivotal role in the integration of somatosensory and auditory informations presenting salient features. This result is in line with a recent functional connectivity study suggesting that the parietal operculum could be a connector hub that links auditory, somatosensory, and motor cortical areas (Tanaka et al., 2018). The connectivity pattern that we obtained is almost similar for both our groups. Checking for significant differences between them, we found a specific connectivity between the tinnitus-related right OP3 seed and the right H-PEEF in the tinnitus group as compared to controls when controlling for the differences in hearing loss and in psychological factors associated with tinnitus, suggesting its relation with the tinnitus percept *per se*. In macaque monkeys, this is the region of the premotor ear-eye field (PEEF) (Lanzilotto et al., 2013), a region that is thought to play an important role in engaging the auditory spatial attention for the purpose of orienting eye and ear towards the sound source. The region found here, posterior to the frontal eye field is probably the human homologue of the PEEF, so an H-PEEF. In macaques, the “premotor ear-eye field” region (PEEF) (Hutchison et al., 2012; Lanzilotto et al., 2013; Lucchetti et al., 2008) involves auditory-motor cells for both eye and ear movements. The human homologue to the macaque PEEF (H-PEEF), for which the connectivity with right OP3 is disturbed in participants with tinnitus is a new finding. It is possible, although speculative, that as in macaques, the H-PEEF might play a premotor role in the control of coordinated ocular movements and tympano-middle-ear movements, pinna movements being vestigial in human. In the right posterior seed, increased connectivity was observed with the Left H-PEEF. Thus, the oculomotor muscles’ proprioceptive system responsible of eyes coordination (Donaldson, 2000) would be disturbed by its potential coupling with an altered middle-ear system. Interestingly, our results could shed light on unexplained observations of poor binocular coordination in subjects with tinnitus. Oculomotor performance during saccades (Lang et al., 2013), fixation, smooth pursuit, optokinetic nystagmus movements and vergence were reduced in tinnitus participants (Josefowicz-Korczynska et al., 2002; Mezzalira et al., 2007; Kapoula et al., 2010; Yang et al., 2010; Lockwood et al., 2001), however signs of vestibular dysfunction were absent. In addition, the literature reports some tinnitus subjects that are able to modulate their tinnitus with eye movements (Coad et al., 2001). These lower performances do not seem to be related to the severity of the handicap because they do not correlate with neuropsychological tests measuring the emotional/attentional status (Josefowicz-Korczynska et al., 2002).

Here we demonstrate that the subregion in the right OP3, already shown to be related to tinnitus (Job et al., 2012; Job et al., 2016), is once again a site of dysfunction in these tinnitus participants. The dysfunction seems focal, because when we further tested a larger seed involving the whole cytoarchitectonic right OP3, we did not elicit any abnormal connectivity from it.

In addition, in the right posterior seed, increased connectivity was observed with the inferior parietal lobule. The inferior parietal lobule is a vast region participating in the integration of sensory information. For instance, the left inferior parietal lobule is involved in proprioceptive illusion (Naito et al., 1999; Kavounoudias et al. 2008; Naito et al. 2017), tactile illusion (Kavounoudias et al. 2008) as well as auditory illusions (Brancucci et al. 2017). These new findings support the view that tinnitus could be a proprioceptive illusion.

Concerning functional connectivity from both Heschl’s gyri, a single differential connectivity was found between our two groups between left Heschl gyrus and the posterior cingulate cortex at the same statistical threshold than for the other opercular seeds. No other significant differential connectivity was found between our two groups, supporting the view that mild tinnitus percept following acoustic trauma is not directly related to modified connectivity with auditory regions. This is complementary to the results from Langers who found that macroscopic tonotopic reorganization in the auditory cortex is not required for the emergence of tinnitus but rather that this reorganization relates to hearing loss (Langers et al., 2012).

## CONCLUSION

In conclusion, our major findings are that: i) in tinnitus participants, there is a specific connectivity between a seed in the right OP3 and a human counterpart to the premotor ear-eye field (H-PEEF), the expression of a crosstalk between ear and eye that may explain disturbed oculomotor performance in tinnitus patients; ii) the small right OP3 seeds also more connected to a region in the inferior parietal lobule in tinnitus patients is confirmed as a key region for mild tinnitus percept, particularly for acoustic trauma tinnitus, being able to integrate somatosensory and auditory salient features. Further investigations will be needed to better understand this disturbed crosstalk between the ear and eye systems and to draw novel conclusions on tinnitus mechanisms.

## ACKNOWLEDGMENTS

The authors are very grateful to Dr F.J. Hemming for proofreading the manuscript, to Dr J.Gauthier, E. Ressiot, H. Michel and C. Klein for selecting tinnitus participants. This work was supported by French government grant number PDH1-SMO-3-0811 (A. Job).

## Author Contributions Statement

Experiment conception and design: A. Job, C. D-M. Experiment execution: A. Job, C. D-M. Data analysis: A. Job, C. D-M, C. J. Discussion about data analysis: A. Jaillard, A. K. Manuscript draft: A. Job, C. D-M, A. Jaillard, A. K, C. J.

## ACRONYMS

CONN: Connectivity toolbox (matlab toolbox)
CSF: Cerebro Spinal Fluid
ENT: Ear, Nose and Throat
EPI: Echo Planar Imaging
FDR: False Discovery Rate
fMRI: functional Magnetic Resonance Imaging
GM: Grey Matter
HL: Hearing levels (dB)
H-PEEF: Human premotor ear-eye field
IPL: Inferior parietal lobule
MNI: Montreal Neurological Institute (brain coordinates system)
MPRAGE: Magnetization Prepared Rapid Acquisition GRE
MR: Magnetic Resonance
OP3: operculum 3
PEEF: premotor ear-eye field
rs-fMRI: resting-state functional Magnetic Resonance Imaging
ROI: Region Of Interest
sd: standard deviation
SPM: Statistical Parametric Mapping
TE: Echo time (ms)
THI: Tinnitus Handicap Inventory
TI: Inversion Time (ms)
TR: Repetition Time (ms)
WM: White Matter

